# Solution and gas-phase modifiers effect on heme proteins environment and conformational space

**DOI:** 10.1101/356444

**Authors:** D. Butcher, J. Miksovska, M. E. Ridgeway, M. A. Park, F. Fernandez-Lima

**Keywords:** trapped ion mobility spectrometry, collision cross section, heme protein, additive, gas-phase modifiers

## Abstract

The molecular environment is known to impact the secondary and tertiary structure of biomolecules, shifting the equilibrium between different conformational and oligomerization states. In the present study, the effect of solution additives and gas-phase modifiers on the molecular environment of two common heme proteins, bovine cytochrome c and equine myoglobin, is investigated as a function of the time after desolvation (e.g., 100 - 500 ms) using trapped ion mobility spectrometry – mass spectrometry. Changes in the mobility profiles are observed depending on the starting solution composition (i.e., in aqueous solution at neutral pH or in the presence of organic content: methanol, acetone, or acetonitrile) depending on the protein. In the presence of gas-phase modifiers (i.e., N_2_ containing methanol, acetone, or acetonitrile), a shift in the mobility profiles driven by the gas-modifier mass and size and changes in the relative abundances and number of IMS bands are observed. We attribute these changes in the mobility profiles in the presence of gas-phase modifiers to a clustering/declustering mechanism by which organic molecules adsorb to the protein ion surface and lower energetic barriers for interconversion between conformational states, thus redefining the free energy landscape and equilibria between conformers. These structural biology experiments open new avenues for manipulation and interrogation of biomolecules in the gas-phase with the potential to emulate a large suite of solution conditions, ultimately including conditions that more accurately reflect a variety of intracellular environments.

## Introduction

Ion mobility spectrometry combined with mass spectrometry (IMS-MS) has increasingly become a complementary or alternative research tool for investigating the conformational space of biomolecules – particularly proteins – under a variety of conditions^1^, including biologically-relevant conditions. For structural applications, IMS-MS studies commonly alter the solution (e.g. salt concentration, pH, temperature) to attempt to emulate a variety of intracellular conditions and sample the protein conformational space (“memory effect” wherein proteins retain a large degree of their solution structure in the gas-phase) ^2^. Solvation of surface residues is an important determinant of protein structure due to charge interactions between surface-exposed residues and nearby water molecules^3^; as a result, protein molecular ions often experience alterations in their structure after desolvation^4^. For example, peptide structures can evolve over time in the gas-phase as indicated by changes in mobility profiles as a function of time^5^. This raises the question of whether it is possible to use bath gas modifiers to change the molecular environment of protein molecular ions during IMS experiments, potentially allowing us to emulate different cell conditions. As we have recently shown, trapped ion mobility spectrometry (TIMS)^6–7^ is particularly well-suited to this approach since ions can be trapped in the gas-phase and exposed to the modified bath gas for variable time periods, typically in the hundreds of milliseconds. In a recent study, we showed the influence of gas-phase modifiers on the conformational space of intrinsically disordered, DNA binding peptide ATHP3 (Lys-Arg-Pro-Arg-Gly-Arg-Pro-Arg-Lys-Trp).^8^

In this study, we use the volatile organic solvents methanol (MeOH), acetone, and acetonitrile (ACN) as solution additives and as gas-phase modifiers to investigate the conformational space of two common heme proteins - bovine cytochrome C (cyt C) and equine myoglobin (Mb) - using TIMS. Our approach differs from other experiments where gas modifiers are used to increase the analytical power of IMS-MS by increasing the size of the collision partner or inducing higher order multi-pole interactions^9^; instead we intend to evaluate whether gas-phase modifiers have the potential to tailor the molecular environment of proteins in the gas-phase to emulate different cell conditions. Results showed that the addition of organic solvents to the starting protein solution can have diverse impacts depending on the protein and the organic solvent (i.e., denaturing or not denaturing the protein). Exposure of protein ions to bath gas modifiers has different effects when compared to the solution additives, from altering the relative abundances of some conformations to creating entirely new IMS bands. This study is the first use of gas-phase modifiers in the TIMS instrument to tune the molecular environment and alter the conformational space of protein molecular ions in the gas-phase.

## Methods and Materials

### Sample preparation

Equine Mb and bovine cyt C (Table S1) were obtained from Sigma Aldrich in lyophilized form and dissolved in Type 1 Ultrapure water. Proteins were exchanged into 10 mM ammonium acetate solution at pH 6.7 immediately before analysis using centrifugal filter units (Amicon Ultra) with a 3 kDa MWCO and diluted to a final concentration of 15 μM.

### Trapped Ion Mobility Spectrometry – Mass Spectrometry Analysis

An overview of the TIMS instrument and its operating principles can be found elsewhere (see Figure S1)^6, 10–11^. In TIMS-MS operation, ions are held stationary against a flow of bath gas by a variable electric field applied along the length of the TIMS tunnel, resulting in ions being simultaneously trapped at different axial positions according to their mobility. The voltage of the tunnel is then ramped to elute ions from the TIMS analyzer, after which they are mass analyzed and detected by a maXis impact Q-TOF MS (Bruker Daltonics Inc, Billerica, MA). Deflector, capillary, and entrance funnel voltage of the TIMS instrument are selected to avoid ion heating/activation prior to TIMS analysis. ^12–13^ The reduced mobility of an ion (Ko) in the TIMS cell is described by:

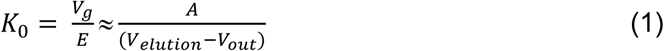

where *v*_*g*_ and *E* are the gas velocity and applied electric field. *V*_*elution*_ and *V*_*out*_ are the elution voltage and base voltage. The constant *A* was determined by calibration with known mobilities of the Tuning Mix calibration standard (G24221A, Agilent Technologies, Santa Clara, CA) in positive ion mode (e.g., *m/z* 622, K_0_ = 1.013 cm^2^ V^−1^ s^−1^ and *m/z* 1222, K_0_ = 0.740 cm^2^ V^−1^ s^−1^)^11^.

Protein sample solutions were directly infused into the inlet of the TIMS instrument via nanoelectrospray ionization (nESI) using laser pulled capillary nESI emitters. HPLC-grade solvents were obtained from Fisher Scientific and used as solution additives and gas-phase modifiers. Solvents were added to sample solutions prior to TIMS-MS analysis. Gas-phase modifiers were added to the TIMS cell by nebulization with the bath gas flow. The gas velocity was kept constant regardless of the bath gas composition. TIMS-MS spectra were analyzed using Compass Data Analysis 5.0 (Bruker Daltonik GmbH) and TIMS Data Viewer 1.4.0.31397 (Bruker Daltonics Inc, Billerica, MA).

## Results and Discussion

Mass spectra show a narrow distribution of charge states – +5 to +8 for cyt C and +6 to +9 for Mb under neutral conditions (pH = 6.9). Notice that, for a solution pH change between from 5.0 to 7.0, the average charge state in solution (determined using PROPKA3^14^ with 2B4Z and 4DC8 reported crystal structures for cyt C^15^ and Mb^16^, respectively) varies from ~12 to ~8 for cyt C and ~9 to ~1 for Mb, indicating that a variety of charge states are accessible in solution between mildly acidic and neutral pH conditions. In most cases, minor differences in the charge state distribution during TIMS-MS were observed for cyt C and Mb as a function of the starting solvent conditions; moreover, a wide charge state distribution was observed in the presence of 50% organic content in the starting solution in the case of Mb in the presence of 50% ACN (Figure 1). Charge state distributions for cyt C and Mb under all solution and modifier conditions are shown in Figure S2. Changes in the total and charge state-derived mobility profiles were studied as a function of the solution starting condition, bath gas composition, and time after desolvation (Figures 2 and 3). The mobility profiles for the low charge states (i.e., +5 - +6 for Cyt C and +7 - +8 for Cyt C) showed a single mobility band, characteristic of native-like solution states that were kinetically trapped during the evaporative cooling of the nESI process, regardless of the starting solvent condition, bath gas composition and time after desolvation. At higher charge states we observe a distribution of unfolding intermediates and fully-unfolded protein, identical to previous ion mobility studies of heme proteins.^17–18^

**Figure 1:**
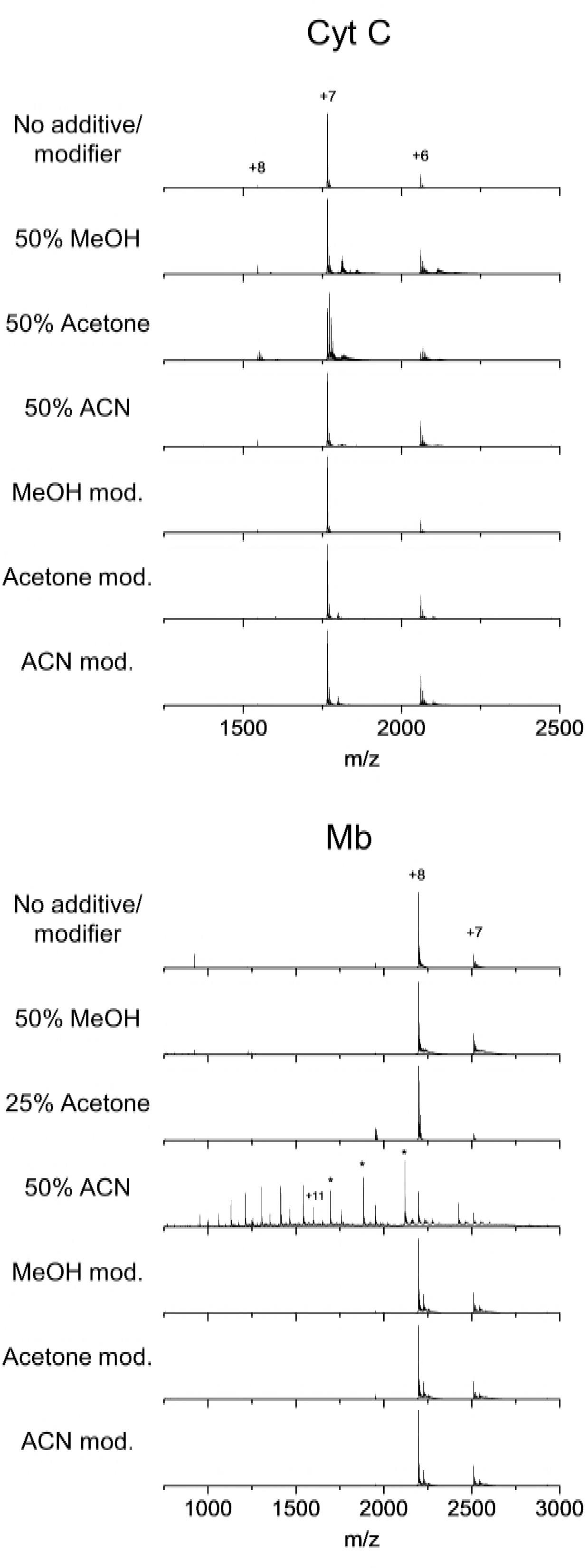
Mass spectra of cyt C (top), and Mb (bottom) with MeOH, acetone, and ACN as solution additives and as gas-phase modifiers.

**Figure 2:**
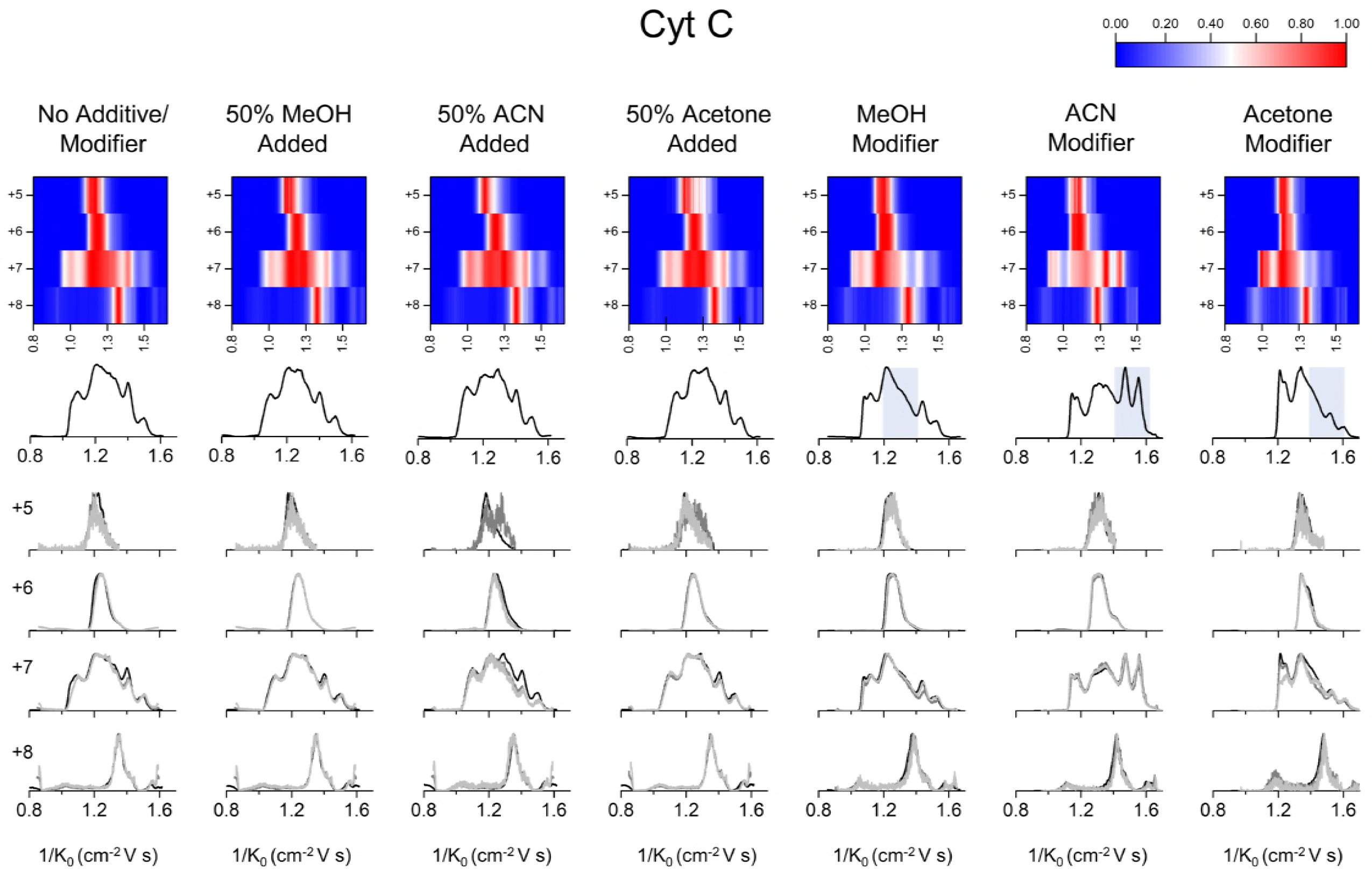
(Top) Heat maps derived from mobility profiles for cyt C showing charge state vs. reduced mobility (1/K0) with normalized intensity across all charge states. (Middle) Combination of highest-intensity mobility profiles for all charge states of cyt C. (Bottom) Mobility profiles determined at 100 ms (black), 300 ms (gray), and 500 ms (light gray) trapping times for all charge states of cyt C.

**Figure 3:**
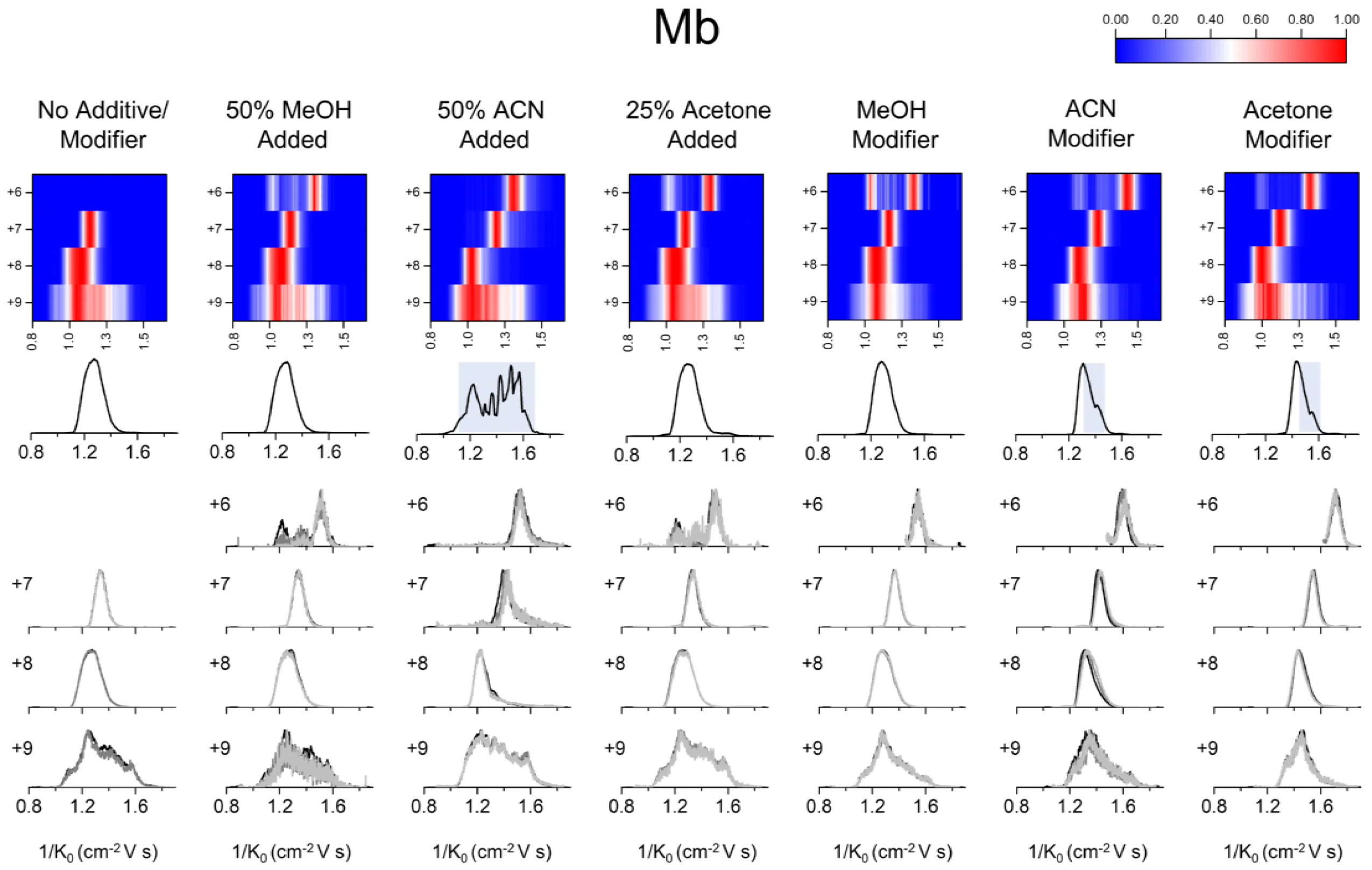
(Top) Heat maps derived from mobility profiles for Mb showing charge state vs. reduced mobility (1/K0) with normalized intensity across all charge states. (Middle) Combination of highest-intensity mobility profiles for all charge states of Mb. (Bottom) Mobility profiles determined at 100 ms (black), 300 ms (gray), and 500 ms (light gray) trapping times for all charge states of Mb.

Mobility profiles of cyt C are not significantly affected by the presence of organic solvents, in good agreement with previous observations (where denaturation was induced by lowering the solution pH). ^17–18^ Mobility profiles of Mb are minimally impacted by the addition of MeOH or acetone, however addition of acetone at 50% v/v causes the protein to precipitate, necessitating the use of a lower volume percentage (25% v/v). 50% ACN causes Mb to unfold to a far greater degree, populating additional charge states (+6 to +17) and conformations, including a single mobility band at the highest charge states (+14 - +17) which likely corresponds to the fully-unfolded protein. Dissociation of heme in the presence of ACN also causes a large increase in the fraction of apo protein (f_apo_) (Table S2) which ranges from 0.64 to 0.86 for charge states +6 to +16 of Mb.

Changes in the mobility profiles of cyt C and Mb are observed using gas-phase modifiers, which are significantly different from those caused by solution additives (Figure 2 and 3, Figure S3 and S4). The mobility shifts correlate across all proteins with the molecular mass and size of the gas-phase modifiers MeOH, ACN and acetone (32, 41 and 58 g mol^−1^, respectively). For example, MeOH as a modifier causes the smallest increase in 1/K_0_, with ACN inducing a larger shift and acetone inducing the largest. While a larger energy transfer (i.e., larger ion effective temperature) is expected with increasing the mass and size of the collision partner, no direct correlation is observed in the IMS profiles; that is, thermally induced changes in the IMS profiles are not observed with the use of modifiers. When the IMS are superimposed, accounting for the mass and size of the gas-phase modifier (Figure S5), a clearer comparison of the modifier effect on the IMS profiles is obtained. While the gas pressure in the TIMS cell is kept constant across the experiments using different gas-phase modifiers, the partial pressure of each modifier in the bath gas differs due to varying vapor pressures of the organic solvents – 16.96, 11.98, and 30.8 kPa for MeOH^19^, ACN^20^ and acetone^21^, respectively (Table S3).

Mobility profiles of cyt C show increased relative abundance for lower-mobility bands in the presence of modifiers, particularly for the +6 - +7 charge states of cyt C and the +7 - +8 charge states of Mb. ACN modifier has a particularly strong impact on the +7 charge state of cyt C, greatly increasing the abundance of the two highest mobility bands which are only weakly observed without the modifier and whose abundance does not increase in the presence of 50% ACN, showing that conformational changes (as reflected by mobility profiles) can be distinct when comparing between exposure of the protein to the organic solvent in the solution-phase or gas-phase.

The changes in the mobility profile observed for cyt C and Mb in the presence of gas-phase modifiers suggests that the protein molecular ions are clustering with the modifier gas molecules while trapped in the TIMS cell. It is known that transient ionic clusters can be formed from modifiers in the bath gas/drift gas and analyte ions generated by electrospray ionization^22^. The nature of the gas-phase modifiers suggests that the main interactions which stabilize site-specific modifier adsorption and cluster formation are based on ion-dipole or dipole-dipole interactions between the modifier and ionized or polar amino acid residues exposed on the protein surface. All gas-phase modifiers used in this study have dipole moments similar to or higher than that of water (i.e., 1.70 D for MeOH, 3.92 D for ACN and 2.69 D for acetone), favoring clustering in the gas-phase. In addition, acetone and ACN have the potential to form stronger ionic interactions than water or MeOH with amino acid residues due to their higher polarity. This hypothesis is in good agreement with previous reports of small peptide clustering in the gas-phase based on a the Langmuir adsorption model^23^ and cluster ion formation during transversal modulation ion mobility^23–24^ and differential mobility spectrometry^25^.

A simplified model based on the transient adsorption of the gas-phase modifiers to the protein molecular ion surface is proposed to explain the changes in the mobility profiles (Figure 4). That is, transient adsorption leads to the observed differences of mobility profiles through disruption of intramolecular contacts, that allows for changes in the energy landscape of the protein favoring a new equilibrium across conformers. Previous studies have shown that upon desolvation, most of the native hydrogen bonds (including those which stabilize secondary structural elements) are maintained and intramolecular contacts are increased overall due to the formation of salt bridges between charged residues on the protein surface^26–27^. Depending on the charge state and protein molecular ion conformation, different reaction pathways can become energetically favorable upon clustering. For example, adsorption of modifiers to charged residues on the protein surface may disrupt surface intramolecular contacts by providing an electrostatic screening effect; this has been observed in the solution-phase^28^ and analogously can be an important determinant of gas-phase protein molecular ion structure. The adsorption of gas-phase modifiers induces a molecular microenvironment at the individual residue level which is more akin to the solution-phase where solvent-exposed residues are solvated by polar molecules which form a solvation shell. This change in molecular environment correlates with differences in relative permittivity at 25 °C between nitrogen gas (~1.0) and the gas-phase modifiers (i.e., 32.7 for methanol^29^, 37.5 for acetonitrile^30^, and 20.7 for acetone^31^). Despite these values being lower than that for water (78.4^32^), residues with adsorbed modifier molecules experience a radically different local electrostatic environment compared to the absence of modifiers. While previous reports using cryogenic IMS have shown the advantages of attaching water molecules to peptides and small proteins as a way to preserve solution-like structures in the gas-phase,^33–34^ the proposed methodology based on TIMS is technically more feasible and opens new avenues for other gas-phase modifiers to reproduce biologically-relevant cell conditions. The main advantages of this approach, when compared to solution studies, are the simultaneous measurement of multiple conformational states and that protein precipitation is no longer a challenge.

**Figure 4:**
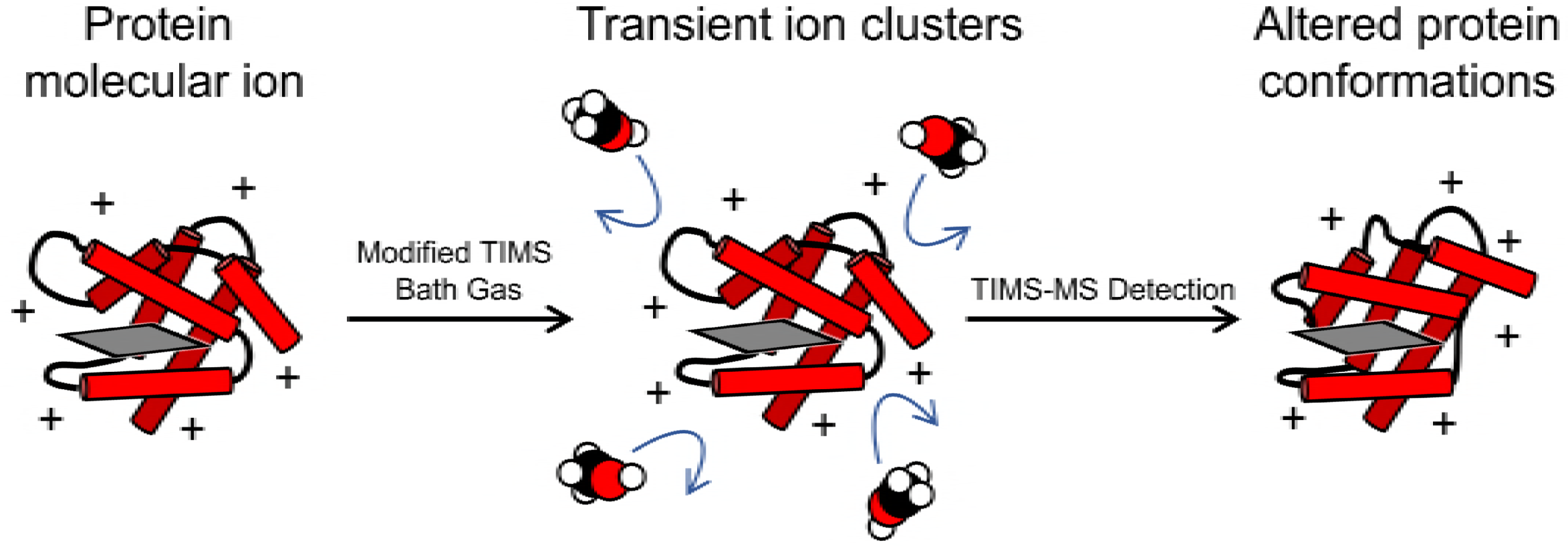
Scheme for adsorption of gas-phase modifiers to the surface of protein molecular ions in the gas-phase. The globin fold of heme proteins is represented by red cylinders while the heme group is shown as a gray square.

## Conclusion

In this study, we observe the impact of the starting solution conditions and the bath gas composition on the conformational space (i.e., mobility profiles) of two common heme proteins. Experimental results showed the presence of memory effects of the starting solution composition on the mobility profiles. We attribute the changes in the number of IMS bands and relative abundances upon addition of gas-phase modifiers to a clustering/declustering mechanism that “solvates” the protein molecular ion through ion-dipole or dipole-dipole interactions, overcoming energetic barriers for conformational interconversion and altering the free energy landscape and equilibrium as a function of the gas-phase modifier. We predict that this process occurs through the solvation of charged residues on the surface of the protein molecular ion by polar modifier molecules which can disrupt intramolecular contacts formed in the gas-phase, as well as providing a charge screening effect. We observe that the addition of gas-phase modifiers to the TIMS bath gas is a completely novel methodology for altering the molecular environment of biomacromolecules in the gas-phase, expanding the possible applications of the trapped ion mobility platform to the study of protein ions in conditions that can emulate a large suite of cell conditions. Future studies will benefit greatly from a comprehensive mechanistic understanding of how clustering in the gas-phase impacts the conformational space of protein molecular ions, allowing investigators to fine-tune the molecular environment to achieve desired outcomes.

## Supporting Information Available

A schematic of the TIMS instrument, charge state distributions for cyt C and Mb with solution additives and gas-phase modifiers, separated mobility profiles for cyt C and Mb with organic solvent additives and gas-phase modifiers for all charge states, shifted mobility profiles for cyt C and Mb in the presence of gas-phase modifiers, sequences and masses of cyt C and Mb, fraction of apoprotein for each state of Mb in the presence of 50% acetonitrile, and physical parameters for organic compounds used as gas-phase modifiers.

## Acknowledgments

This work was supported by the National Science Foundation Division of Chemistry, under CAREER award CHE-1654274, with co-funding from the Division of Molecular and Cellular Biosciences to F.F.-L. We would like to thank Antonija Tangar and Riupeng Lei for contribution of protein samples.

## Declaration of Competing Interests

The authors declare no competing interests.

